# Biophysical Principles of Lineage Factor PU.1 Binding Revealed by NextPBMs

**DOI:** 10.1101/328625

**Authors:** Nima Mohaghegh, David Bray, Jessica Keenan, Ashley Penvose, Kellen K. Andrilenas, Vijendra Ramlall, Trevor Siggers

## Abstract

Determining the biophysical principles that shape transcription factor (TF) binding in a cell-specific manner is key to quantitative models of gene expression. High-throughput (HT) *in vitro* methods measuring protein-DNA binding are invaluable for relating TF binding affinity to genome-wide binding; however, the impact of cell-specific post-translational modifications (PTMs) and cofactors are not routinely assessed. To address these limitations, we describe a new HT approach, called nextPBMs (<u>n</u>uclear <u>ext</u>ract <u>p</u>rotein-<u>b</u>inding <u>m</u>icroarrays), to characterize TF binding that accounts for PTMs and endogenous cofactors. We use nextPBMs to examine the DNA binding of the lineage factor PU.1/Spi1 and IRF8 in human monocytes. We identify two binding modes for PU.1 in monocytes – autonomous binding unaffected by PTMs and cooperative binding with IRF8, and identify a single cooperative mode for IRF8. We characterize the DNA binding of PU.1:IRF8 complexes, and show how nextPBMs can be used to discover cell-specific cofactors and characterize TF cooperativity at single-nucleotide resolution. We show that chromatin state and cofactors both influence the affinity requirements for PU.1 binding sites. Furthermore, we find that the influences of cooperative (IRF8) and collaborative (C/EBPα) cofactors on PU.1-binding-site affinity are independent and additive.

## INTRODUCTION

Determining the biophysical factors that dictate where a transcription factor (TF) binds throughout the genome remains a challenge. Primary determinants of TF binding are protein-DNA affinity and cooperative binding with other TFs (Siggers and Gordân, 2013; Slattery et al., 2014). Various high-throughput (HT) *in vitro* techniques are currently available to characterize the DNA binding of TFs (reviewed in (Andrilenas et al., 2015; Slattery et al., 2014)), and cooperative TF complexes (Jolma et al., 2015; Siggers et al., 2011; Slattery et al., 2011). These approaches have provided rich datasets of TF-DNA binding information and provided a tremendous resources for analyzing genomic data (Weirauch et al., 2014). Current HT methods use either purified or *in vitro* produced protein(Badis et al., 2009; Berger et al., 2006a; Slattery et al., 2011), or tagged protein overexpressed in cells (e.g., HEK293) (Fang et al., 2012; Jolma et al., 2013). Consequently, these approaches do not assay the impact of cell-specific post-translational modifications (PTMs), which are known to have diverse effects on TF binding and function(Filtz et al., 2014; Tootle and Rebay, 2005). Furthermore, these approaches do not implicitly account for the impact of cell-specific cofactors that can bind cooperatively with a target TF to affect its DNA binding. To address these limitations, we describe a new HT approach, called nextPBMs (<u>n</u>uclear <u>ext</u>ract PBMs), to characterize the binding of a TF that accounts for the effect of cell-specific PTMs and cofactors. We apply the nextPBM approach to examine the biophysical determinants of binding for PU.1/Spi1 protein in human monocytes.

The ETS family protein PU.1 is a master regulator of the myeloid lineage (Nerlov and Graf, 1998; Rosenbauer and Tenen, 2007; Scott et al., 1994) and functions to establish localized histone modifications that define the monocyte/macrophage-specific enhancer repertoire (Barozzi et al., 2014; Ghisletti et al., 2010; Heinz et al., 2010). PU.1 can function autonomously to establish enhancers (Ghisletti et al., 2010) and macrophage-specific gene expression (Feng et al., 2008); however, it also functions with other TFs (Heinz et al., 2010; Langlais et al., 2016; Mancino et al., 2015; J. A. Zhang et al., 2012). PU.1 binds DNA cooperatively with IRF8 to ETS-IRF composite elements (EICEs) (Eklund et al., 1998; Meraro et al., 1999; Rehli et al., 2000), and the absence of IRF8 reduces PU.1 binding to EICEs in macrophages (Mancino et al., 2015). PU.1 can also collaborate with C/EBPα to bind chromatin and establish macrophage-specific genes expression (Feng et al., 2008; Heinz et al., 2010; Laiosa et al., 2006; Xie et al., 2004); however, PU.1 does not bind DNA cooperatively with C/EBPα via direct protein-protein interactions, and their collaboration is likely through mutual effects on repressive chromatin environments. In addition to cofactors, PU.1 binding is also impacted by nucleosomes, and is inhibited at DNA sequences with high nucleosome occupancy (Barozzi et al., 2014). Therefore, genome-wide PU.1 binding is dependent on DNA-binding affinity, nucleosomes, and interactions with cofactors.

To better understand how DNA binding affinity relates to genome-wide PU.1 occupancy, we used nextPBMs to measure PU.1 binding to thousands of binding sites identified by ChIP-seq in human THP-1 monocytes. Previous work examining the relation between PU.1-DNA binding affinity and *in vivo* occupancy found that binding sites tend to be lower affinity at loci co-occupied with other proteins (Pham et al., 2013), demonstrating that co-occupancy with other proteins alters the affinity requirements for PU.1 binding. However, these studies did not examine the impact of co-occupancy with the major monocytes cofactors IRF8 or C/EBPα. Additionally, previous studies have relied on position-weight matrices (PWMs) to model PU.1 binding affinity (Barozzi et al., 2014; Pham et al., 2013); however, PWMs do not always reliably predict the impact of DNA shape (Barozzi et al., 2014; Zhou et al., 2015), extended flanking DNA (Gordân et al., 2013; Le et al., 2018), and do not implicitly account for PTMs or cooperative binding with cofactors. Here, we have taken a more direct approach and measured PU.1 binding to genome-derived sites with their native flanking sequences; we have used nextPBMs to account for monocyte-specific PTMs and cofactors; and we have directly examined the impact of interactions with cooperative (IRF8) or collaborative (C/EBPα) cofactors on the affinity requirements for *in vivo* PU.1 binding.

## RESULTS

### Genome-wide Binding for PU.1, C/EBPα and IRF8 in human monocytes

To define the PU.1 binding landscape in human monocytes, and co-occupancy with cofactors, we performed ChIP-seq on PU.1, C/EBPα and IRF8 in resting THP-1 cells (**Methods**). We observed widespread binding for each factor and significant overlap in their binding profiles (Figure 1A), consistent with previous studies (Ghisletti et al., 2010; Heinz et al., 2010; Langlais et al., 2016; Mancino et al., 2015). We identified 47,799 PU.1, 26,648 C/EBPα and 2588 IRF8 binding loci (i.e., ChIP-seq peaks) in resting THP-1 cells. The number of PU.1 binding sites is consistent with numbers reported for human peripheral blood monocytes (Pham et al., 2012)and mouse macrophages (Ghisletti et al., 2010; Heinz et al., 2010; Mancino et al., 2015). Critically, we observed near complete overlap of IRF8 binding sites (95%) with PU.1 binding sites, supporting the model that *in vivo* IRF8 must bind as a complex with PU.1.

**Figure 1.**
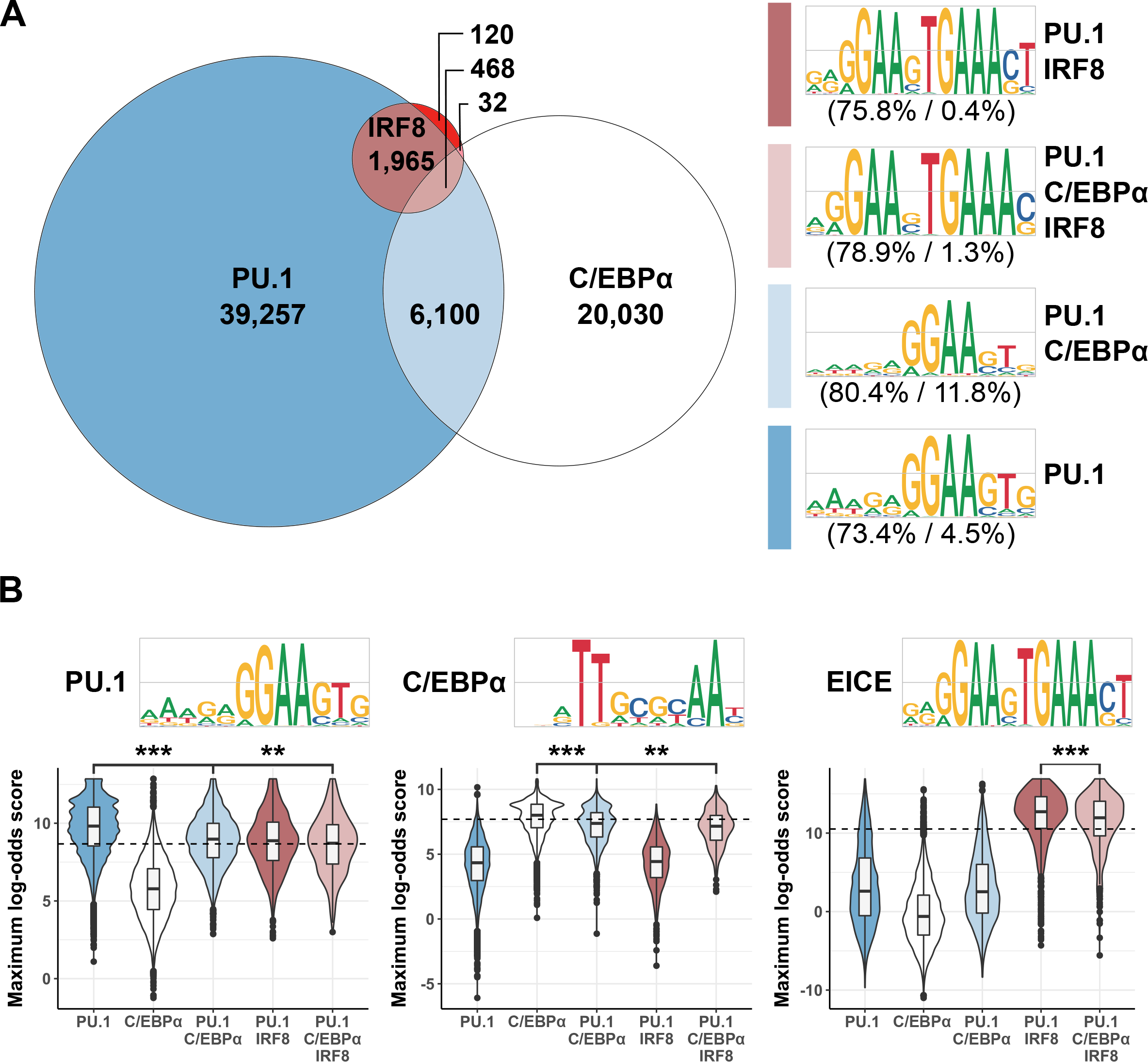
Genome-wide Binding for PU.1, C/EBPα and IRF8 in human monocytes. **(A)** Overlap of genome-wide ChIP-seq peaks discovered for PU.1, C/EBPα, and IRF8. *De novo* motifs discovered within each PU.1-containing intersection are shown on the right. Numbers in brackets indicate the percentage of peaks containing the *de novo* motif (left) compared to background (right). Grey bars in the motif logos represent bit values of 0 (bottom), 1 (middle), and 2 (top). When a ChIP-seq peak overlapped multiple peaks from another experiment we aggregated them into a single overlapping region. **(B)** Distributions of motif scores obtained for ChIP-seq peaks categories described in (A). ChIP-seq peaks were scanned with each motif and assigned the maximum log-odds score (see **Methods**).

To characterize the DNA sequence motifs that define PU.1, C/EBPα and IRF8 binding we performed *de novo* motif analysis on defined subsets of the bound genomic loci (**Methods**). Binding motifs determined for genomic loci bound by PU.1 alone or with other factors agree well with known motifs (Figure 1A). At loci bound by PU.1 alone we identify a canonical PU.1 binding motif. At loci shared with IRF8 (or IRF8 and C/EBPα) the dominant motif is the EICE bound cooperatively by IRF8 and PU.1. At loci bound with C/EBPα we identify a PU.1 motif, supporting the idea that collaboration between these factors is not via direct cooperative DNA binding but rather through synergistic effects on chromatin (Feng et al., 2008; Ghisletti et al., 2010). We find that the PU.1 motif identified on loci co-occuied with C/EBPα is slightly more degenerate than for PU.1 alone, suggesting that PU.1 binding sequences may be lower affinity at loci shared with C/EBPα (discussed more below).

To determine the specificity of TF motifs for their respective *in* vivo binding profiles we scored bound regions using the individual TF motifs (Figure 1B). For both PU.1 and C/EBPα we observe that their binding motif is highly predictive of their ChIP-seq peaks. However, for both TFs, motif scores are lower for loci bound with the other factors. For IRF8, we find that the EICE motif scores are much higher for IRF8 ChIP-seq peaks than for peaks from other factors. These analyses demonstrate that motifs identified for each TF are specific for their genomic binding loci, and that TF binding at loci co-occupied with either a collaborating or a cooperative co-factor can be lower scoring.

### Characterizing PU.1 binding in monocytes using NextPBMs

To characterize the role of binding affinity on PU.1 occupancy we sought to directly measure PU.1 binding to thousands of binding sites found in the PU.1 ChIP-seq peaks. As cell-specific PTMs and cofactors can perturb TF binding affinity, we developed a HT approach – nextPBMs (<u>**n**</u>uclear <u>**ext**</u>ract **PBMs**) – to measure TF-DNA binding directly from nuclear extracts (Figure 2A). PBMs are double-stranded DNA microarrays that allow *in vitro* measurement of protein binding to tens of thousands of unique DNA sequences (Berger et al., 2006b). NextPBMs extend the PBM methodology by using nuclear extracts in place of purified, IVT, or over-expressed proteins (**Methods and Materials**). As proteins are not epitope tagged, microarray-bound proteins are probed with native primary antibodies followed by fluorescently labeled secondary antibodies. Critically, in nextPBMs, the assayed proteins have all endogenous PTMs and are in the presence of all cell-type-specific cofactor proteins.

**Figure 2.**
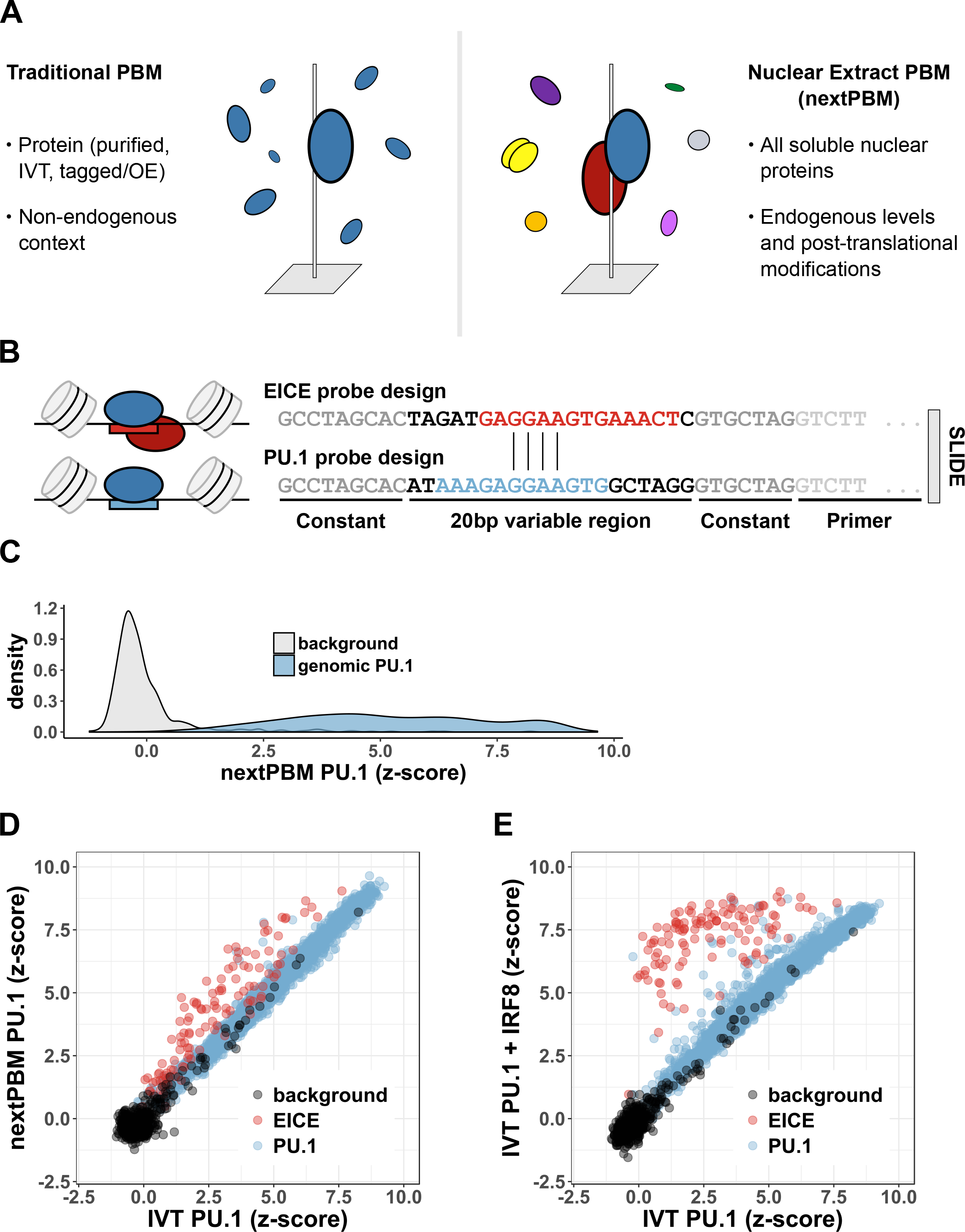
Characterizing PU.1 binding in monocytes using nextPBMs. **(A)** Schematic of traditional PBM and nextPBM experiments. **(B)** Microarray probe schematic. EICE is highlighted in red. Canonical PU.1 motif is highlighted in blue. Genomic nucleotides flanking the motifs are indicated in black. For each probe, 20-bp genomic variable regions are embedded within constant flanking DNA. 24 nucleotide primer sequence is included on each probe and used to double-strand the probes (see **Methods**). **(C)** Density of PU.1 nextPBMΔz-scores obtained at random background probes (n = 500) and at genomic PU.1 binding sites (n = 2,615).
**(D)** Comparison of PU.1 bindingΔz-scores determined by IVT-PU.1 PBM and PU.1 nextPBM. Binding is shown to 116 EICEs (red), 2,499 canonical PU.1 binding sites (blue), and 500 random background probes (black). **(E)** Comparison of PU.1 bindingΔz-scores determined by IVT-PU.1 PBM and IVT PU.1+IVT IRF8 PBM. Binding sites are as in (D).

To analyze the binding of PU.1 from human monocytes, we performed nextPBMs using THP-1 nuclear extracts and monitored the binding of PU.1 to select binding sites from our PU.1 ChIP-seq data. Binding was assayed to 2,615 genomic PU.1 sites, including composite PU.1:IRF8 EICEs, to survey PU.1 binding to a diverse set of genomic loci bound by different cofactors. Binding was assayed to 20-bp-long sites to account for the impact of genomic sequence context (i.e., flanking DNA)(Figure 2B). PU.1 bound to genome-derived sites with much higher affinity (i.e., higher PBM z-scores) than to 500 randomly selected background sites (Figure 2C), demonstrating that there is sufficient PU.1 protein in nuclear extracts to effectively assay protein-DNA binding in our assay.

To determine how cofactors or PTMs affect PU.1 binding in a nuclear extract context, we compared the nextPBM PU.1 binding data with binding of PU.1 generated by *in vitro* translation (IVT) (Figure 2D). We observe excellent correlation for the majority of PU.1 sites between nuclear extract PU.1 and IVT PU.1 binding data (Figure 2D, highlighted in blue); however nuclear extract PU.1 binding was enhanced to the EICEs (Figure 2D, highlighted in red) found in genomic regions co-occupied by PU.1 and IRF8. The enhanced binding seen by nextPBM suggests cooperative binding of PU.1 to these sites with IRF8 present in the nuclear extract. To determine whether the enhanced binding to EICEs is similar to that expected using only PU.1 and IRF8, we performed a PBM with IVT PU.1 and IVT IRF8 (Figure 2E). As expected, EICEs are preferentially bound by PU.1 in the presence of IRF8, while the bulk of the remaining canonical PU.1 sites are unaffected. The 2,615 PU.1 binding sites in our nextPBM sample approximately 5% of the PU.1 ChIP-seq peaks. For this diverse sample of PU.1 sites, enhanced binding of PU.1 is observed only to EICEs; therefore, we reason that IRF8 is likely the only factor in resting human monocytes binds cooperatively with PU.1. Our results demonstrate that in human monocytes PU.1 binds in two modes – autonomously to canonical PU.1 sites in a manner that is unaffected by PTMs, and cooperatively with IRF8 to EICEs.

### Sequence determinants of two PU.1 binding modes in monocytes

To analyze the sequence determinants of PU.1 binding we queried the base preferences along select binding sites and generated DNA-binding logos using a single-nucleotide variant (SNV) approach (**Materials and Methods)**(Andrilenas et al., 2018). Briefly, we measure the binding of PU.1 to 20 bp-long *seed* sequences and all 60 SNV sequences; logos were then generated directly from binding scores to each SNV sequence (Figure 3A-D, **Methods and Materials**). This approach defines a logo for *a single seed sequence* and allows us to probe the nature of binding to individual cooperatively or non-cooperatively bound sites.

**Figure 3.**
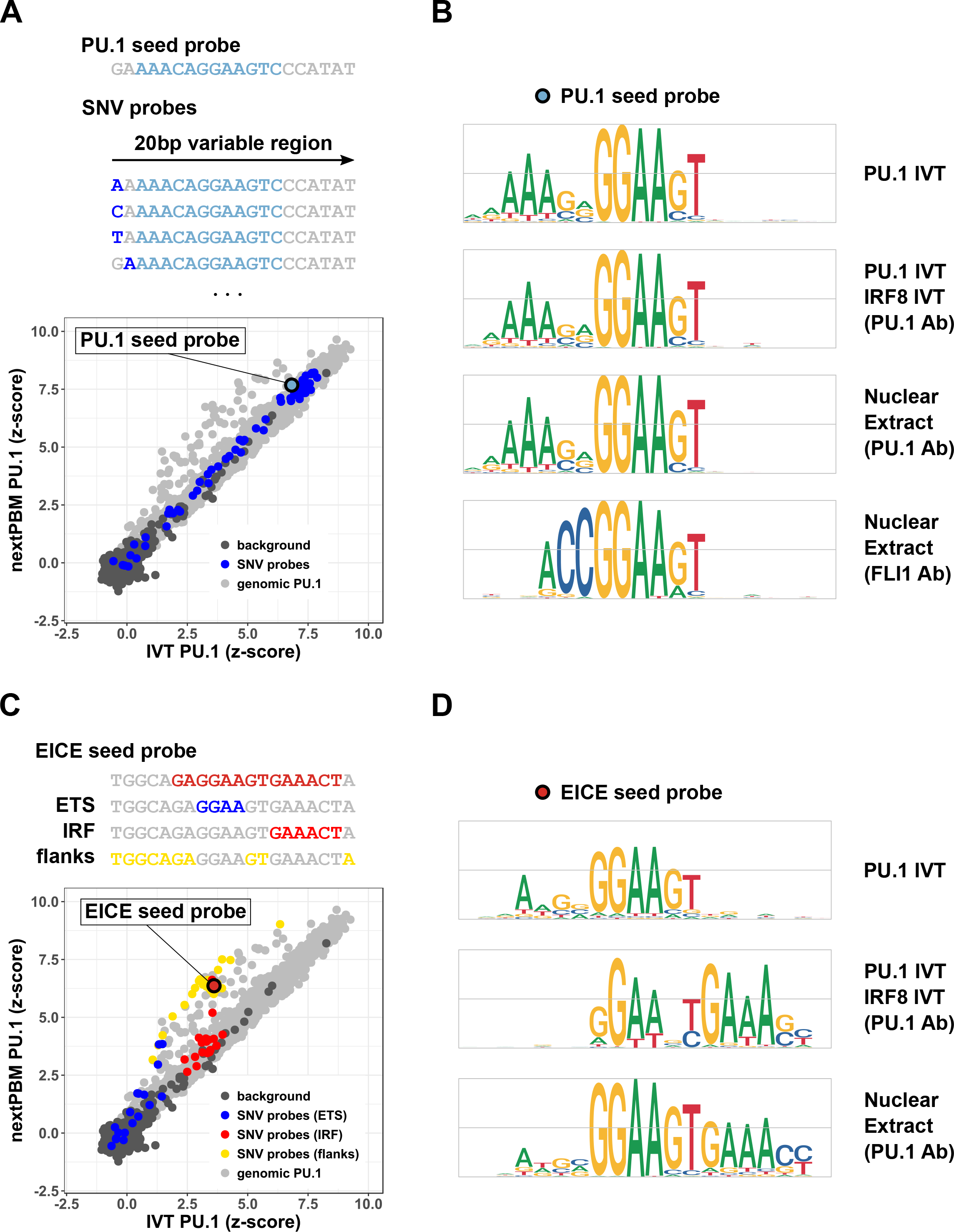
Sequence determinants of two PU.1 binding modes in monocytes. **(A)** Top – schematic of single-nucleotide variant (SNV) probes generated relative to a canonical genomic PU.1 seed sequence. Canonical PU.1 binding site within the 20-bp variable region is highlighted in light blue. Bases that correspond to SNVs are shown in dark blue. Bottom - scatterplot of PU.1 bindingΔz-scores as in Figure 2D. SNVs of the indicated canonical PU.1 seed probe are shown in dark blue. **(B)** Motif logos constructed using the PU.1 seed probe and its corresponding SNV probes (see **Methods**) for different nextPBM and PBM experiments. **(C)** Top – schematic of EICE seed probe and bases comprising individual sub-element. Bottom – scatterplot of PU.1 bindingΔz-scores as in Figure 2D. Highlighted is binding to SNV probes containing variations in either the ETS core site (blue), IRF core site (red), or flanking and linker bases (yellow). **(D)** Motif logos constructed as in (B) using the indicated EICE seed probe and its corresponding SNV probes. Grey bars in the motif logos represent bit values of 0 (bottom), 1 (middle), and 2 (top).

We identified distinct motifs for PU.1 based on cooperatively or non-cooperatively bound seed sequences (Figure 3A-D). PU.1 binding to SNVs of a non-cooperatively bound seed sequence was highly correlated for IVT and nuclear extract conditions (Figure 3A) and lead to the established PU.1 motif under both conditions (Figure 3B). An identical motif is found for IVT PU.1 in the presence of IVT IRF8. These results show that the presence of IRF8 has no effect on PU.1 binding to canonical PU.1 sites. In contrast, binding to SNVs of a cooperatively bound seed sequence produced an EICE motif, with an IRF8 binding site (5’-GAAACT-3’) adjacent to the PU.1 core site (5’-GGAA-3’) (Figure 3D). For this same seed sequence the IVT PU.1 experiment produces a canonical PU.1 motif, while the IVT PU.1 plus IVT IRF8 experiment reveals the EICE motif (Figure 3D). These results demonstrate that the extended EICE motif is due to the presence of IRF8. To ensure that cooperative binding with IRF8 was specific to PU.1, and not a feature of any ETS factor protein in our assay, we used nextPBMs to examine the binding of FLI1. FLI1 is an ETS factor expressed at high levels in THP-1 cells. Binding of FLI1 to the non-cooperatively bound seed sequence yielded a canonical FLI1 logo(Wei et al., 2010) that is distinct from the PU.1 logo(Figure 3D), while binding to the cooperatively bound seed sequence was not significant in our assay(data not shown). In summary, using a single nextPBM experiment, we recovered the two known PU.1 binding logos that describe the sequence preferences for cooperative and non-cooperative binding modes.

The ability to identify the full EICE motif by only monitoring the binding of PU.1 suggests that we can use nextPBM experiments to both identify and dissect the biophysical determinants of cooperative TF binding. To further probe the sequence determinants of cooperative PU.1:IRF8 binding, we looked at the effect of SNVs in different regions of the EICE (Figure 3C). SNVs in the 5’-GGAA-3’ PU.1 core site abrogate PU.1 binding for both IVT and extract samples, demonstrating the absolute requirement of the ETS site (highlighted in blue). SNVs in the 5’-GAAACT-3’ IRF8 site affect the cooperative binding without affecting the binding of PU.1 alone (highlighted in red). This suggests that cooperative binding with IRF8 does not alter the binding specificity of PU.1; rather IRF8 simply binds to an adjacent site and enhances PU.1 binding. This is further supported by SNVs in the flanks and linker bases that affect PU.1:IRF8 complex affinity but do not abrogate the cooperative interactions (i.e., all points are above the diagonal) (highlighted in yellow). To examine the determinants of PU.1:IRF8 binding more broadly, we examined PU.1 binding to a more diverse set of 60 EICE elements from PU.1:IRF8 co-occupied genomic loci (Figure 4A). EICE sites are all bound well above background and are enhanced in nuclear extracts compared to IVT PU.1 due to cooperative binding with IRF8. Mutating the ETS site in the EICEs completely abrogates binding of PU.1, whereas abrogating the IRF8 site only affects the cooperative enhancement. Finally, we asked whether the apposition of a strong IRF8 binding site next to a weak PU.1 motif would be sufficient to enhance PU.1 binding. We generated 199 synthetic EICEs by combining low-affinity PU.1 sites with a consensus IRF8 site, and assayed PU.1 binding by nextPBM. An adjacent IRF8 site greatly enhanced PU.1 binding to all sites in the presence of the nuclear extract but not for IVT PU.1 (Figure 4D). Strikingly, these synthetic EICEs showed even greater cooperativity than the native EICEs with stronger PU.1 binding sites. These results demonstrate that from nextPBM experiments we can identify and characterize composite cooperative binding elements at nucleotide resolution.

**Figure 4.**
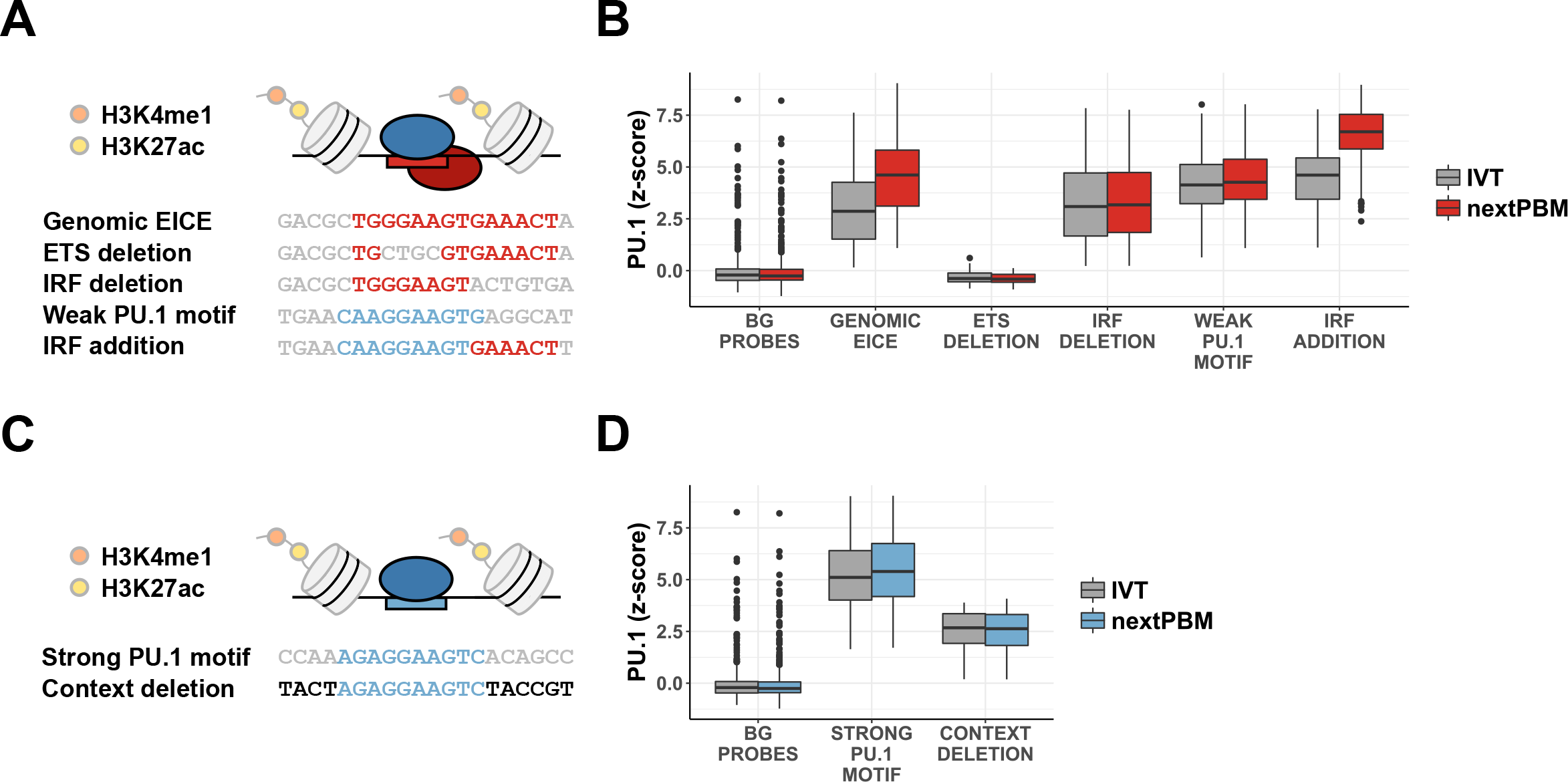
PU.1 binding to synthetic mutations of genomic sites. **(A)** Schematic showing representative probe sequences and corresponding mutated elements. For each genomic EICE from active enhancers (n = 60), there is a corresponding probe with the ETS and IRF half-sites independently deleted. For each canonical PU.1 probe with a weak motif (n = 199), there is a corresponding probe with an IRF half-site added. BG – background. **(B)** Distributions of PU.1 bindingΔz-scores for DNA probe groups in (A) in nuclear extract (nextPBM) versus IVT. **(C)** Schematic showing a representative genomic PU.1 site from an active enhancer with a strong motif (highlighted in light blue) and its genomic context beyond the 10bp core motif (light blue) deleted (black). For each of these genomic PU.1 sites (n = 539), there is a corresponding probe with the context deletion applied. **(D)** Distributions of PU.1 bindingΔz-scores for DNA probes groups in (C) in nextPBM versus IVT.

The impact of DNA sequence flanking a TF binding site (i.e., flanking DNA) can have considerable effects on protein-DNA binding that may not be accurately capture by PWMs (Gordân et al., 2013; Le et al., 2018). Our PBM-determined PU.1 binding logos have a stretch of Ade bases (i.e., A-tract) 5-prime to the core 5’-GGAA-3’ that has been reported for PU.1 (Barozzi et al., 2014; Jolma et al., 2013; Pham et al., 2013) and that is also present in our ChIP-derived motifs (Figure 1). This A-tract demonstrates that PU.1 binds to a longer binding site than has been reported for other ETS factors (Wei et al., 2010) and may be strongly influenced by flanking DNA sequence. Furthermore, recent studies have suggested that PU.1 may actually bind as a dimer(Esaki et al., 2017), suggesting even greater importance for an extended DNA site. To test this we altered the flanking sequence of 539 non-cooperative PU.1 sites and measured the impact on PU.1 binding (Figure 4D). We observed that the alteration of the flanking DNA had a significant impact on the PU.1 binding, decreasing the average z-score by almost 2. These results highlight the importance of flanking DNA bases for accurate measurements of PU.1 binding.

### Sequence determinants of IRF8 binding in monocytes

IRF8 requires cooperative interactions with PU.1 for high-affinity binding to EICEs (Eklund et al., 1998; Meraro et al., 1999; Rehli et al., 2000). Consistent with this requirement for PU.1, most IRF8 occupied regions are co-occupied with PU.1 (95 %) (Figure 1). To biophysically examine this IRF8 requirement, we measured IRF8 binding by nextPBM to PU.1 sites and EICEs indentified in our ChIP-seq data. We find that IRF8 binds almost exclusively to the EICE elements (Figure 5A). Furthermore, we note that the nextPBM z-scores are much higher for IRF8 than for PU.1 due to very low binding of IRF8 to background sequences, which is consistent with its reported inability to bind DNA on its own. To examine the sequence determinants of IRF8 binding, we generated a binding logo using the same EICE seed sequence examined for PU.1 in Figure 3. The logo matches the known EICE motif, consistent with a requirement for PU.1 cooperativity (Figure 5). This result also demonstrates that we recover the same binding logo by nextPBM by probing either member of the cooperative TF complex. To more fully test the requirement for cooperative binding with PU.1, we examined the impact on IRF8 binding to 60 EICEs for which we mutated either the PU.1 ETS site or the IRF8 site. Binding of IRF8 is completely abrogated with mutations to either the ETS sites or the IRF8 site, demonstrating that IRF8 binding is dependent on cooperativity(Figure 5B). Finally, to determine whether cooperativitiy might be dependent on the PU.1 binding site sequence, we examined IRF8 binding to synthetic EICEs formed by apposing an IRF8 site next to a range of 199 weak PU.1. IRF8 bound strongly to these synthetic EICEs, and at levels higher than seen for the native EICEs (Figure 5B). These results demonstrate that IRF8 requires an adjacent PU.1 site (with a range of possible affinities) to bind EICEs in monocytes.

**Figure 5.**
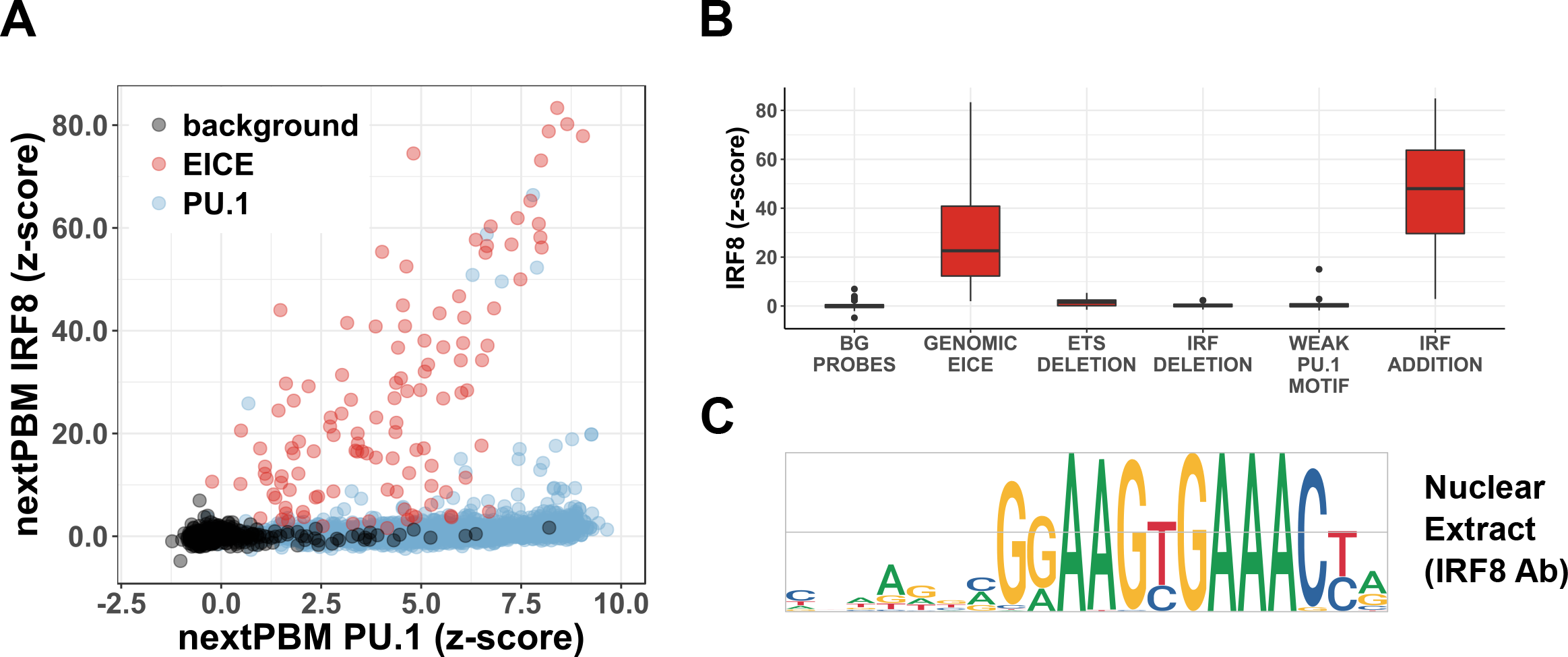
Sequence determinants of IRF8 binding in monocytes. **(A)** Scatterplot of IRF8 bindingΔz-scores in nuclear extract compared to *in vitro* translated (IVT) at genomic EICEs (n = 116, red), canonical PU.1 binding sites (n = 2,499, blue), and random background probes (n = 500, black). **(B)** Distributions of IRF8 bindingΔz-scores for DNA probe groups from Figure 4 in nuclear extract (nextPBM) versus IVT. **(C)** Motif logo constructed as in Figure 3 using IRF8 bindingΔz-scores from IRF8 nextPBM experiment to the indicated EICE see probe. Grey bars in the motif logo represent bit values of 0 (bottom), 1 (middle), and 2 (top).

### Role of cofactors on PU.1 binding affinity

To examine how PU.1 binding affinity is affected by co-occupancy with either a collaborating (C/EBPα) or cooperative-binding (IRF8) cofactor, we compared PU.1 binding to sites from regions co-occupied by PU.1 and the cofactors C/EBPα and IRF8. To control for chromatin state, we performed genome-wide ChIP-seq for histone 3 lysine 4 mono-methylation (H3K4me) and H3K27 acetylation (H3K27ac) that define poised (H3K4me only) and active (H3K4me and H3K27ac) enhancer states (Creyghton et al., 2010; Heintzman et al., 2007), and analyzed PU.1 binding to sites identified only in active enhancer regions. To control for potential indirect interactions, we analyzed only ChIP-positive regions in which a binding motif for the TF was present (**Methods**). For each genomic region, PU.1 binding was measured to the PU.1 site with the highest PWM score.

To examine the impact of IRF8 on PU.1 affinity, we compared PU.1 binding to EICEs (from regions co-occupied with IRF8) with sites found in regions occupied PU.1 alone (Figure 6A). The binding of IVT PU.1 (i.e., absence of any cooperativity) is significantly lower affinity to EICEs than to sites from PU.1-only-bound regions (Δz-score ~ 2.0, p-value < 0.001). However, as expected, this PU.1 binding difference is much less significant in the nextPBM experiment where PU.1 is bound cooperatively with IRF8 (Δz-score ~ 0.5, p-value < 0.001). It is possible that low affinity PU.1 binding to EICEs is due to the sequence constraints imposed by presence of the IRF8 binding site. However, we showed that binding of PU.1 alone is unaffected by base changes to the IRF8 binding site (Figure 3D, 4B). Furthermore, we showed that juxtaposing an IRF8 site and a weak PU.1 site does not lower binding of PU.1 alone, but rather creates a highly cooperative EICE (Figure 4B). These results demonstrate that cooperative binding with IRF8 allows the EICEs to be lower affinity PU.1 sites.

**Figure 6.**
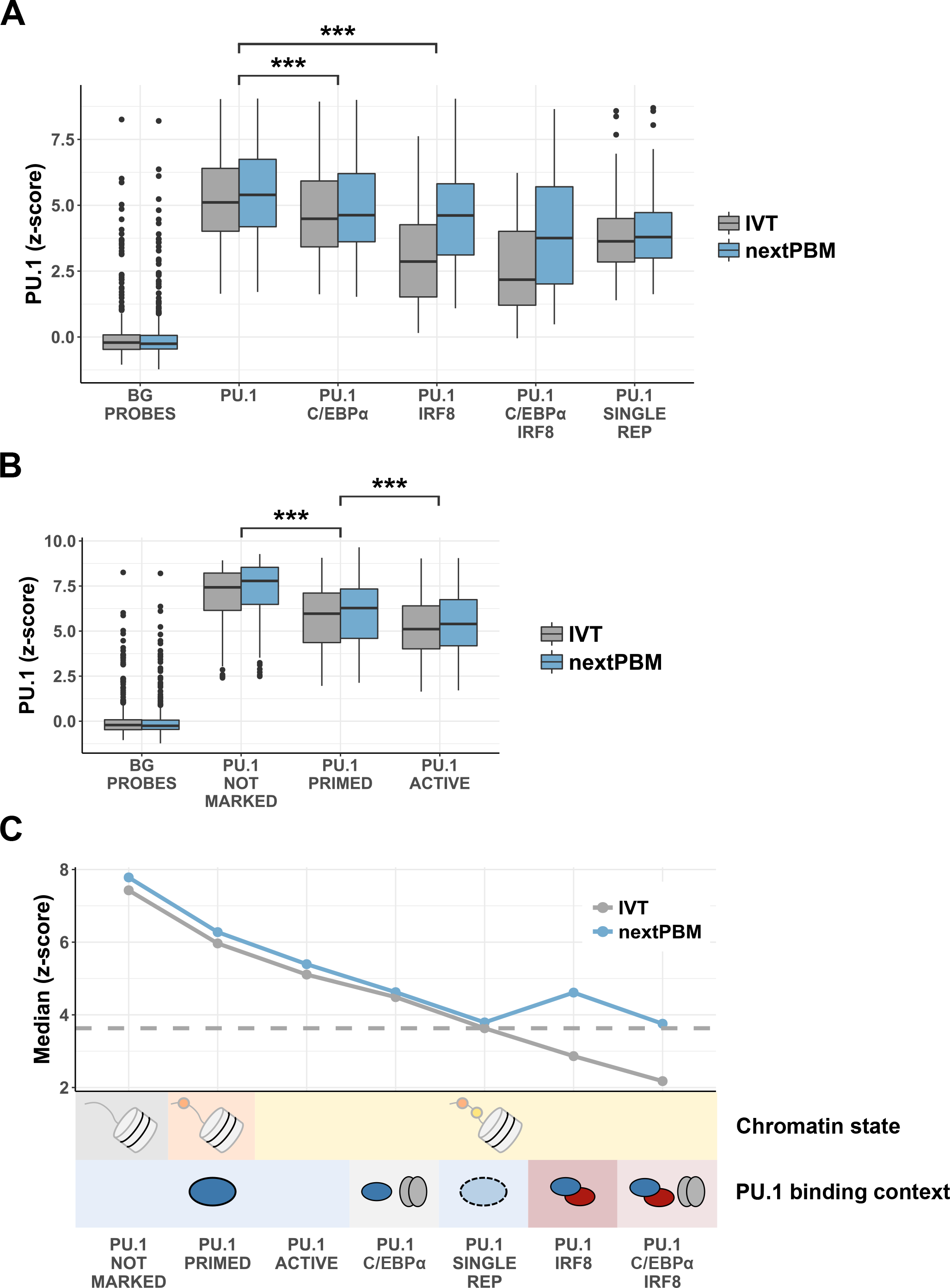
PU.1 binding in different chromatin and cofactor contexts. **(A)** Distributions of PU.1 binding z-scores for DNA probe categories defined by ChIP-seq co-occupancy with cofactors. All PU.1 sites are from ChIP-seq peaks that overlap with active enhancers. ‘SINGLE REP’ designates a category of probes from PU.1 ChIP-seq peaks that were discovered in a single experiment but could not be duplicated. **(B)** Distributions of PU.1 binding z-scores for DNA probe categories consisting of genomic PU.1 sites located in different enhancer states. ‘NOT MARKED’ indicates the absence of H3K4me or H3K27ac histone marks. **(C)** Schematic summarizing the trends observed in (A) and (B) for PU.1 binding site affinity in different chromatin states and cofactor contexts.

To examine the impact of C/EBPα co-occupancy on PU.1 affinity, we compared PU.1 binding to sites in regions occupied by PU.1 alone or co-occupied with C/EBPα (Figure 6A). PU.1 binding is also lower affinity for sites from C/EBPα co-bound regions (Δz-score ~ 0.5, p-value < 0.001), although the magnitude is less than seen for IRF8 co-bound regions, and this trend is true for both nextPBM and IVT PU.1 experiments. Unexpectedly, the same trend of lower-affinity PU.1 binding with C/EBPα co-occupancy is also observed for composite EICEs (Δz-score ~ 0.5, p-value = 0.051). This trend was further supported by our initial motif-based analysis of the full ChIP-seq dataset (Figure 1B, p-value < 0.001). Therefore, for both modes of PU.1 binding, co-occupancy with C/EBPα allows PU.1 binding to be lower affinity. Combining the impact of both cooperativity and collaboration, we see that the lowest affinity PU.1 binding sites are those in regions bound by both IRF8 and C/EBPα (Δz-score ~ 2.5). These observations suggest that the impacts of cooperative and collaborative cofactors on the affinity requirements for PU.1 are independent and additive.

### Role of affinity in ChIP-seq reproducibility

To determine whether lower affinity PU.1 binding observed for some PU.1 occupied regions might be related to binding dynamics or assay reproducibility, we examined regions identified as PU.1 bound in only a single ChIP-seq replicate experiment (‘single-replicate regions’). For many single-replicate regions, PU.1 binding sites were readily identified. Binding of IVT PU.1 to these single-replicate sites was lower affinity than for reproducible PU.1 sites (Figure 6A). This suggests that ChIP reproducibility is related to PU.1 binding affinity. Based on the binding affinity to single-replicate regions, we defined an affinity threshold to demarcate reproducible PU.1 binding (Figure 6A, dashed line). We find that the majority of PU.1 loci have sites with affinity above this threshold, and that PU.1 binding to EICEs is above the threshold only in the presence of IRF8. Therefore, despite the complications of chromatin structure, we find that PU.1 binding affinity measured by nextPBM can broadly discriminate reproducible PU.1 binding sites.

### Role of affinity in PU.1 binding to distinct chromatin environments

The observation that PU.1 binding is lower affinity at regions co-occupied by C/EBPα suggests that chromatin environment impacts the affinity of PU.1 sites. Previous studies have demonstrated that PWM motif scores for PU.1 binding sites are highest in inaccessible chromatin regions defined by low DNAse I accessibility (Pham et al., 2013), suggesting that the chromatin accessibility relates to PU.1 binding site affinity. We compared the PU.1 binding to sites identified in gene-distal regions classified by distinct enhancer states: primed, active, or unmarked (no H3K4me or H3K27ac marks) (Figure 6B). To control for the effect of C/EBPα and IRF8, we considered only sites from PU.1-only occupied loci. PU.1 binding affinity shows a clear trend with enhancer state, with the lowest affinity binding in active enhancers and highest affinity in unmarked loci. High-affinity PU.1 binding to unmarked loci is in agreement with previous studies (Pham et al., 2013) and suggests that PU.1 occupancy to unmarked and less biophysically accessible chromatin regions requires high-affinity sites. Primed enhancers can be viewed as enhancers that may become active in a different monocyte cell state; however, we should then expect PU.1 binding-site affinity to be the same for active and primed enhancer states. Therefore, the moderate but significant (p-value < 0.001) difference in PU.1 affinity between these classes suggests that either PU.1 binding to these regions is opportunistic (J. A. Zhang et al., 2012) or many of these primed enhancers are not PU.1-dependent enhancers in different cellular conditions. Finally, the observation that PU.1 binding is lowest affinity at active enhancers suggests that functional PU.1 sites are not the highest affinity, and that genome-wide analyses of the highest affinity TF sites may be enriched for non-functional binding.

## DISCUSSION

Relating biophysical principles of TF binding to *in vivo* occupancy is critical to a mechanistic view of gene regulation. Cell-specific PTMs(Filtz et al., 2014; Tootle and Rebay, 2005) and cofactors(Garvie and Wolberger, 2001; Siggers and Gordân, 2013) can affect TF binding, but are not implicitly accounted for in current HT methods for analyzing protein-DNA binding. Here we describe the nextPBM methodology for the HT characterization of protein-DNA binding that accounts for cell-specific PTMs and cofactors. We demonstrate that nextPBMs can be used quantitatively characterize the binding of TFs directly from cell nuclear extracts, and that by comparing binding of purified/IVT and nuclear extract samples we have a genome-scale approach to discover cooperatively bound DNA sequences. Integrating ChIP-seq (to identify genomic regions of interest) and nextPBMs (to characterize protein binding) provides a powerful approach to examine TF binding that accounts for cell-specific differences.

As all nuclear proteins are present in a nextPBM experiment, the method assays, in parallel, the DNA-binding of a TF in potentially multiple distinct protein complexes. We use a SNV approach (Andrilenas et al., 2018) to generate binding logos and thereby probe the biophysical configuration of the complexes bound to different DNA sequences. Applying this approach to sites bound by PU.1 cooperatively and non-cooperatively allowed us to recover the two known PU.1 binding logos from a single experiment, and to quantitatively characterize the sequence determinants of cooperative PU.1:IRF8 binding at nucleotide resolution. Importantly, by assaying either PU.1 or IRF8 we identified the composite EICE binding logo, which provides information about the identity of the other partner (i.e., a prediction if the partner is unknown). Therefore, coupling nextPBMs and the SNV approach provides a general approach to discover and characterize cooperative TF binding complexes.

Analyzing the binding of PU.1 from human monocytes, we identified two binding modes – autonomous binding that is unaffected by PTMs, and cooperative binding with IRF8. In contrast, we found a single cooperative binding mode for IRF8 to the composite EICE. The cooperative binding requirement for IRF8 agrees with biochemical studies (Eklund et al., 1998; Meraro et al., 1999; Rehli et al., 2000), and agrees with ChIP-seq analyses that show co-binding of PU.1 at essentially all IRF8-bound sites in mouse macrophages (Langlais et al., 2016; Mancino et al., 2015) and human monocytes (Figure 1). In other cell types and cell conditions, PU.1 and IRF8 are known to form TF complexes with other cofactors. For example, IRF8 is proposed to bind cooperatively with IRF1 in LPS-and IFNγ-stimulated mouse macrophages (Langlais et al., 2016; Mancino et al., 2015), and in HEK293 cells can bind with AP-1 family members JunB and BATF(Glasmacher et al., 2012). In B cells, PU.1 can bind cooperatively with IRF4 (Brass et al., 1999). These studies highlight that cooperative binding to composite sites is integral to the cell-specific function of PU.1 and IRF8. NextPBMs provide a HT approach to discover and characterize new cooperative interactions for these regulators in additional cell types and conditions.

Investigating the role of affinity in genome-wide PU.1 binding revealed a striking relationship between PU.1 binding affinity and both chromatin state and cofactor occupancy (Figure 6C). Analyzing chromatin state, we found a decreasing trend for PU.1 binding affinity with the presence of enhancer marks - the highest affinity PU.1 binding occurs in regions not marked as enhancers, followed by primed enhancers, then active enhancers. This agrees with previous studies that found PU.1 PWM scores are highest in inaccessible chromatin regions defined by low DNAse I accessibility (Pham et al., 2013). These results suggest that in repressive chromatin environments only the highest affinity PU.1 sites can be bound, while at active enhancers the DNA is more biophysically accessible and PU.1 requires lower affinity sites to bind. Analyzing PU.1 binding with cofactors, we found that binding with C/EBPα (a collaborative cofactor) can be lower affinity than for PU.1 alone (Δz-score ~ 0.5), and that PU.1 binding with IRF8 (a cooperative factor) can be even lower (Δz-score ~ 2). Unexpectedly, we found that thebiophysically distinct influences of these two cofactors on PU.1 affinity are independent and additive, as the lowest PU.1 affinity binding sites are in regions co-occupied by both cofactors, and their combined impact on PU.1 affinity was additive (Δz-score ~ 2.5). These results suggest that distinct biophysical mechanisms can independently contribute to lowering the evolutionary constraints on TF binding site affinity required for *in vivo* occupancy.

The observed trends observed for PU.1 affinity with chromatin state and co-factor binding reveal important concepts related to both the biophysical principles of TF binding and potential biases in computational genomic analyses. First, functional PU.1 sites are not among the highest affinity sites, therefore, computational analyses focused on high-scoring sites for PU.1 (and TFs more generally) will be biased to non-functional binding loci. Second, cooperatively bound PU.1 sites are among the lowest affinity PU.1 sites and score poorly by standard PWMs making them easy to miss in genomic analyses; however, they are almost exclusively in active enhancers and, therefore, likely functionally relevant. We propose that the combined ChIP-seq/nextPBM approach presented here provides a useful new approach to examine the biophysical principles of TFs in gene regulation that accounts for these observed affinity trends and addresses complications from cell-specific PTMs and cofactors.

## ACKNOWLEDGEMENTS

We thank J. Fuxman Bass for helpful comments on the manuscript. This work was supported by a NIH R01 grant (R01A116829 to T.S.)

## AUTHOR CONTRIBUTIONS

Experiments, N.M., T.S.; Computational analyses, D. B. and T.S; Technology development, N.M, D.B., J.K, A.P., K.K.A, V.R., T.S; Writing, N.M., D.B, T.S.; Funding, T.S.

---

**Supplementary File 1. NextPBM/PBM Data**

**Supplementary File 2. Transcription Factor Motifs and Thresholds**

**Supplementary File 3. NextPBM/PBM Experimental Details**

## METHODS

### Cells, Protein Samples, Antibodies

<u>***Cells***</u> - THP-1 cells were purchased from ATCC (cat # TIB-202). <u>***Protein samples***</u> – *In vitro* transcription/translation (IVT) samples of PU.1 and IRF8 (full-length, untagged) were generated using 1-Step Human Coupled IVT Kit – DNA (Thermo Fisher Scientific Cat# 88881) and following the provider’s instructions for cloning the corresponding cDNAs into the IVT compatible plasmid (provided in the kit) and setting up reactions for protein expression. We confirmed each protein-expression by western analysis. <u>***Antibodies***</u> – PU.1 (Santa Cruz sc-352x, used for ChIP and nextPBM); C/EBPα (Santa Cruz sc-61x, used for ChIP); IRF8 (Santa Cruz sc-6058x, used for ChIP and nextPBM); human histone 3 lysine 4 mono methylation (H3K4me) (Abcam ab8895, used for ChIP); histone 3 lysine 27 acetylation (H3K27ac) (Abcam ab177178, used for ChIP); alexa488-conjugated anti-goat (Life Technologies A1105, used for nextPBM); alexa488-conjugated anti-rabbit (Life Technologies A11034, used for nextPBM); and FLI (ABclonal A5644, used for nextPBM) was a gift from ABclonal.

### Nuclear Extracts

5 × 10^6^ THP-1 cells pelleted at 500xg for 5 minutes at 4°◻ in a 15 ml conical tube. The pellet resuspended and washed twice with PBS. Cell pellet was resuspended in 1 ml of ‘low-salt buffer’ (10 mM HEPES (pH 7.9), 1.5 mM MgCl2, 10 mM KCl plus 1 μl protease inhibitor cocktail (Sigma-Aldrich, cat # P8340) and incubated for 10 minutes on ice. 50 μl of 5% IGEPAL (Sigma-Aldrich, cat # I8896) was added to the cell suspension followed by a harsh vortex for 10 seconds. Released nuclei were immediately pelleted at 750xg for 5 minutes at 4°◻. The supernatant was transferred to another tube and saved as the ‘cytosolic fraction’. To wash the remaining cytosolic proteins from the surface of the nuclear pellet, 100 μl of the low-salt buffer was gently pipetted onto the side of the tube and allowed to wash the pellet, but making sure to not disrupt the pellet. This wash was then gently transferred to the cytosolic fraction without dislodging the nuclear pellet. 200 μl of ‘high-salt buffer’ (20 mM HEPES (pH 7.9), 25% Glycerol, 1.5 mM MgCl2, 0.2 mM EDTA, 420 mM NaCl plus 1 μl protease inhibitor cocktail) was pipetted on the pellet and the tube went through a vigorous vortex for 30 seconds followed by nutation at 4°◻ for 1 hour. The nuclei were pelleted at 4°◻ for 20 minutes at 21,000xg. The supernatant was transferred into another tube as the nuclear soluble protein fraction. Final nuclear extract samples used in nextPBM assays were 9.6 mg/ml.

### ChIP-Seq

4 × 10^7^ THP-1 cells per ChIP-seq experiment were grown in RPMI medium with 10% fetal bovine serum (FBS). Soluble-chromatin was prepared according to previously described protocols (Lee et al., 2006)with some modifications. Briefly, cells were crosslinked with 1% formaldehyde (final concentration) (Fisher Scientific, cat # F79-500) for 10 minutes at room temperature with gentle shaking. Crosslinking was stopped by adding glycine solution in PBS (125 mM final concentration). Fixed cells were pelleted at 800xg for 5 minutes at 4°◻ and washed twice with 10 ml of cold PBS in a 15 ml conical tube and pelleted at 800xg for 5 minutes at 4°◻. Washed cell pellet was resuspended in 10 ml of Lysis Buffer 1(Lee et al., 2006), nutated for 10 minutes at 4°◻, and pelleted at 2000xg for 5 minutes at 4°◻. The same procedure was repeated with Lysis Buffer 2(Lee et al., 2006) at room temperature followed by pelleting at 2000xg for 5 minutes at 4°◻. To release nuclei from hard-to-disrupt THP-1 membranes, cells were resuspended in 10 ml of Lysis Buffer 3(Lee et al., 2006) and were shaken vigorously (225 rpm) at room temperature for 30 minutes. Cells then were passed through an 18-gauge needle (VWR, cat # BD305195) for 25 times using a 10ml syringe. Nuclei were pelleted at 3000xg for 20 minutes at 4°◻ and resuspended in 500 μl of Lysis Buffer 3 and then transferred into a 1.5 ml microfuge tube placed in Benchtop 1.5 ml Tube Cooler (Active Motif, cat # 53076). The nuclei were sonicated using Active Motif Q120AM sonicator with a 3.2 mm Probe (Active motif cat # 53053) at 25% amplitude for 15 minutes with 20 seconds ON and 30 seconds OFF cycles (45 cycles total). Cell-debris was pelleted at 21,000xg for 30 minutes at 4°◻. 50 μl of the combined soluble-chromatin was saved to be used as the input DNA upon reverse-crosslinking. For immunoprecipitation, 500 μl of the soluble chromatin was mixed with 30 μg of either PU.1, C/EBPα, H3K4me or H3K27ac antibodies (60 μg of IRF8 antibody was mixed with 1 ml of the soluble chromatin), and a tubes were rotated at 25 rpm for one hour at 4°◻ using HulaMixer (ThermoFisher Scientific cat # 15920). 125 μl of the protein A Dynabead slurry (ThermoFisher Scientific cat # 10001D) per each rabbit antibody (PU.1. C/EBPα, H3K4me or H3K27ac), and 250 μl of the protein G Dynabead slurry (ThermoFisher Scientific cat # 10003D) for the goat-IRF8 antibody, were transferred into 1.5 ml microfuges and placed on DynaMag magnet (ThermoFisher Scientific, cat # 12321D) until all beads collected on the side of tubes. The solution was gently aspirated off from each tube and then the rack was removed from the magnet and beads were resuspended in 1 ml of the Lysis Buffer 3 with several gentle inversions and then the rack was put back on the magnet and the buffer was aspirated off after all beads were collected on the side of the tube. Each bead then resuspended in 50 μl of Lysis Buffer 3 and were added to the corresponding tube and returned to HulaMixer to rotate at 35 rpm for over-night at 4°◻. Dynabeads were collected with placing the tubes in the magnet. Beads were washed for 6 times with 1 ml of the Lysis Buffer 3 and two times with 1 ml of the Wash Buffer (RIPA). All ChIP samples along with the 50 μl of the soluble-chromatin were reverse-crosslinked by adding 200 μl of the Elution buffer and 3 μl of 20 mg/ml Proteinase K (ThermoFisher Scientific, cat # AM2546) and incubated at 65°◻ for over-night. Beads were collected by magnets and the solutions were transferred into a new 1.5 microfuges containing 1 μl of 10 mg/ml RNase A (ThermoFisher Scientific, cat # EN0531) and left at room temperature for an hour. The ChIP and input DNA were purified using QIAquick PCR Purification Kit (QIAGEN, cat # 28104) and eluted in 50 μl of 50°◻ Nuclease-Free Water (Thermo Fisher Scientific, AM9932). The concentration and size distribution of the ChIP-DNA samples were defined using Agilent 2100 Bioanalyser. DNA libraries were prepared using NEBNext Ultr II DNA Library Prep kit (NEB, E7645S) following the provider’s instruction manual. Amplified libraries were Bioanalyzed again to check the size selection efficiency and to define the concentrations of libraries before preparing the library pool involving the same molarity of each library and sequenced by Illumina NextSeq 500.

### ChIP-seq analysis

ChIP-seq reads were aligned to the human reference genome (hg19) using Bowtie2(Langmead and Salzberg, 2012). Aligned reads were filtered for high quality and uniquely mappable reads (MAPQ > 30) using samtools(Li et al., 2009). Peak calling for TFs was performed using MACSv2 (Y. Zhang et al., 2008)with relaxed parameters on single experiments (p-value < 0.01) and experiments were filtered using the irreproducible discovery rate (IDR < 0.05) across biological duplicates(Landt et al., n.d.). Peak calling for histone marks was performed using MACSv2 (Y. Zhang et al., 2008)with relaxed parameters on single experiments (p-value < 0.01) and experiments were filtered requiring identification in both biological duplicates (i.e., IDR was not used for histone marks analysis). Peaks were further filtered if they occurred in the ENCODE consortium blacklisted regions. Peak intersections were computed using bedtools(Quinlan and Hall, 2010) by first merging the peaks from all transcription factor ChIP-seq experiments into continuous genomic loci and identifying which TF(s) contained a peak within this union set. Raw and processed ChIP-seq data is available in the NCBI Geo database (Accession #XXX).

### Motif discovery and scoring

*De novo* motifs within peak sets were discovered using HOMER(Heinz et al., 2010) and subsequently used for motif scoring across all peaks. Log-odds scoring thresholds determined by HOMER against a set of random background sequences were used as significance thresholds for motif scanning. Motif scans (Figure 1) on individual peaks were performed using a custom R script that implements the same scoring scheme as HOMER and reports the maximum log-odds score in each peak. Uniform background probability for each nucleotide (0.25) at each position was used for log-odds scoring. Log-odds score boxplots were generated using the ggplot2 R package(Wickham, 2016). Motif logos were generated using the ggseqlogo R package(Wagih, 2017). Motifs and thresholds used for ChIP-seq analysis and PBM microarray design are provided (**Supplementary File 2**).

### PBM design

PBM experiments were performed using custom-designed microarrays (Agilent Technologies Inc. AMADID 085106, 8x60K format). 2,615 PU.1 binding sites identified in ChIP-seq peaks were extracted from the genome as 20-bp genomic fragments and placed into a fixed position in the PBM probe sequence (Figure 1). For each unique probe sequence, 5 replicate probes were included in each orientation (10 probes per unique site). For select genomic seed sequences, 60 matching SNV probes were included to assay all single-nucleotide variants at the 20 positions of the binding site (Figure 3). All SNV sites were also included with 5 replicates and in each orientation (10 probes per unqiue SNV site). Probes for assaying binding site ablations and synthetic EICE sites were similarly included with 10 probes per unique DNA site. <u>***Selection of binding sites from ChIP-seq data***</u> Binding sites were only included from PU.1 ChIP-seq peaks demonstrating high reproducibility across biological duplicates (IDR < 0.01), with the exception of probes included specifically to assay binding to ‘single-replicate’ regions. PU.1 ChIP-seq sites were categorized based on their log-odds motif score, proximity to co-factors, and enhancer state. PU.1 binding sites were selected from the PU.1 ChIP-seq peaks containing exactly one significant PU.1 site (see Motif analysis above). For some genomic loci we identified no significant PU.1 site and, therefore, used the PU.1 site with maximum log-odds score (Figure 2 **and 3**). EICE sites were selected from PU.1-IRF8 co-occupied regions containing exactly one EICE site (see Motif analysis above). Co-occupancy PU.1 with co-factors (C/EBPα and/or IRF8) was determined if a highly reproducible ChIP-seq peak (IDR < 0.01) for each factor over-lapped by at least one base. A PU.1 ChIP-seq peak peak was annotated as ‘PU.1-alone’ if it was located greater than 200 bases away from the nearest cofactor ChIP-seq peak (in all experiments, including duplicates, with peaks called using relaxed parameters as detailed above). Enhancer states were annotated using histone modification ChIP-seq data from biological duplicates and publically available mRNA-seq data for THP-1 monocytes (GEO accession GSM927668). PU.1 sites were annotated as active if they occurred within 200 bases of the nearest H3K4me1 and H3K27ac peaks, and if the nearest gene was located between 2-500kb away and expressed above the median RPKM value. PU.1 sites were annotated as primed if they occurred within 200 bases of the nearest H3K4me1 peak only, and if the nearest gene was located between 2-500kb away and expressed below the median RPKM value. A full list of DNA probes used, the genomic location from which they originated (where applicable), their corresponding probe category and additional annotation can be found in the supplemental data (**Supplementary File 1**).

### NextPBM and PBM experiments and analysis

Microarray DNA double stranding and basic PBM protocols are as previously described (Berger and Bulyk, 2009; Siggers et al., 2015). All wash steps were carried out in coplin jars on an orbital shaker at 125 rpm. Double-stranded DNA microarrays were first pre-washed in PBS containing 0.01% Triton X-100 (5 min), rinsed in a PBS bath, and then blocked with 2% milk in PBS for 1 hour. Following the blocking step, arrays were washed in PBS containing 0.1% Tween-20 (5 min), then in PBS containing 0.01% Triton X-100 (2 min), and finally briefly rinsed in a PBS bath. <u>***Protein binding***</u> - Arrays were then incubated with the protein sample (IVT protein or THP-1 nuclear lysate, details in **Supplementary File 3**) for one hour in a binding reaction buffer containing: 2% milk (final concentration); 20 mM Hepes buffer, pH 7.9; 100 mM NaCl; 1 mM DTT; 0.2 mg/mL BSA; 0.02% Triton X-100; and 0.4 mg/mL salmon testes DNA (Sigma D7656). <u>***Primary antibody***</u> - After protein incubation, microarrays were washed with PBS containing 0.5% Tween-20 (3 min), then in PBS containing 0.01% Triton X-100 (2 min), followed by a brief PBS rinse. Microarrays were then incubated with 10 μg/mL of primary antibody (see **Supplementary File 3**) in 2% milk in PBS (20 min). <u>***Secondary antibody***</u> – After primary antibody incubtion, microarrays were washed with PBS containing 0.5% Tween-20 (3 min), then in PBS containing 0.01% Triton X-100 (2 min), followed by a brief PBS rinse. Microarrays were then incubated with 7.5 μg/mL of alexa488-conjugated secondary antibody (see **Supplementary File 3**) in 2% milk in PBS (20 min). Excess antibody was removed by washing with PBS containing 0.05% Tween-20 (3 min), then PBS (2 min). <u>***PBM data analysis***</u> - Microarrays were scanned with a GenePix 4400A scanner and fluorescence was quantified using GenePix Pro 7.2. Exported data were normalized using MicroArray LINEar Regression (Berger et al., 2006b). Microarray probe sequences are provided (**Supplementary File 1**). PBM data analysis and SNV approach to logo generation is a previously described (Andrilenas et al., 2018). Processed PBM z-score data is available in the supplementary data (**Supplementary File 1**), and all raw PBM data in the NCBI Geo database (Accession #XXX). Scatterplots and boxplots were generated using the ggplot2 R package (Wickham, 2016). Motif logos were generated using the ggseqlogo R package(Wagih, 2017). The significance of PU.1 binding affinity and motif scores between groups was calculated using the Wilcoxon-Mann Whitney test implemented in R.

